# The pathogenic mutant vimentin L387P disrupts endoplasmic reticulum organization and proteostasis

**DOI:** 10.64898/2025.12.07.692853

**Authors:** Elena Sala Lara, Dolores Pérez-Sala, Alma E. Martínez

**Affiliations:** Department of Cellular and Molecular Biosciences, Centro de Investigaciones Biológicas Margarita Salas, C.S.I.C., 28040 Madrid, Spain

**Author notes:** To whom correspondence should be addressed at: Centro de Investigaciones Biológicas Margarita Salas, C.S.I.C., Ramiro de Maeztu, 9, 28040 Madrid, Spain.

**Keywords:** vimentin L387P, progeroid syndrome, ERQC, ERAD, endoplasmic reticulum, proteostasis, ER dysfunction

## Abstract

The organization and dynamic regulation of the vimentin intermediate filament network are essential for diverse cellular processes, including cell division, cytoskeletal crosstalk, and organelle positioning. In particular, the vimentin network is required to maintain a compact perinuclear endoplasmic reticulum (ER) that supports the ER quality-control compartment. Here, we show that the vimentin variant c.1160T>C (L387P), identified in a progeroid syndrome, disrupts ER organization and function. In SW13/cl.2 cells, vimentin L387P forms aggregates containing cisternal-like structures of predominant perinuclear localization that associate with pronounced nuclear distortion. These aggregates sequester chaperones, the SEL1L/HRD1 complex, and the 20S proteasomal subunit, while excluding RNF26 and VCP/p97. This extensive redistribution of ER-associated degradation components coincides with impaired proteostasis and reduced free Ca^2+^ in intra-organelle compartments, indicating compromised ER functionality. Altogether, these findings suggest that the L387P mutation drives extensive ER disorganization and highlight the importance of an intact vimentin network for ER homeostasis.

## Introduction

Vimentin is a cytoskeletal protein belonging to the type III intermediate filament (IF) class that predominantly assembles into a dynamic and resilient filamentous network spanning the cytoplasm. In addition, vimentin can be detected at the cell surface or in the extracellular space, either in soluble or vesicle-bound forms, or as oligomeric structures [1]. The vimentin monomer comprises a central αhelical rod domain flanked by intrinsically disordered head and tail regions. Parallel association of monomers yields coiled-coil dimers, which further assemble in an antiparallel configuration to form tetramers, the fundamental precursors of filament assembly. Subsequent lateral alignment of tetramers gives rise to unit-length filaments (ULFs), short oligomeric intermediates that later elongate longitudinally to generate mature intermediate filaments [2,3]. The assembly of vimentin filaments is a highly dynamic and reversible process, enabling subunit exchange, lateral association and elongation, and disassembly in response to cellular cues [4]. This is tightly regulated by posttranslational modifications, including phosphorylation, ubiquitination, or oxidation, which modulate filament organization and turnover during processes such as mitosis, migration, and cellular stress responses [5–7].

The vimentin network confers mechanical stability and spatial organization to the cell, and is essential for the proper positioning and morphology of organelles such as the nucleus, mitochondria, Golgi apparatus, lysosomes, and endoplasmic reticulum (ER) [8–13]. Focusing on the ER, recent studies have revealed that vimentin is essential for maintaining the structural integrity of the perinuclear ER. Specifically, vimentin acts as an anchoring scaffold for perinuclear ER sheets, supporting ER qualitycontrol (ERQC), which is critical for protein degradation, suggesting that vimentin is a key determinant of not only ER morphology and dynamics but also ER function [12].

The ER regulates many cellular pathways, including protein and lipid synthesis, protein quality control, and calcium storage [14]. Upon exposure to cellular insults, including oxidative stress, proteins may undergo posttranslational modifications that can potentially compromise their folding, leading to the accumulation of misfolded species and possibly triggering ER stress [15,16]. In turn, the unfolded protein response (UPR) is activated to restore ER homeostasis by increasing its folding capacity and directing terminally misfolded proteins to the ER-associated degradation (ERAD) pathway. ERAD constitutes a key quality control mechanism that prevents the accumulation of aberrant proteins within the ER by recognizing, ubiquitinating, and retrotranslocating misfolded or unassembled proteins to the cytoplasm for proteasomal degradation [17,18], thereby preserving ER function and overall cellular proteostasis. Disruption of this finely tuned balance has been implicated in numerous human pathologies including metabolic, neurodegenerative, cardiovascular, or age-associated diseases, as well as cancer [19–21]. Notably, vimentin wild-type (wt) has been shown to interact closely with several ERAD members. Such interactions appear to serve a dual purpose. Vimentin associates with E3 ubiquitin ligases, such as RNF208 or TRIM56 or their adaptors, namely gigaxonin, to regulate its own turnover, being targeted for degradation [22–25]. In addition, vimentin contributes to stabilize specific ERAD components, supporting their proper function through its association with the E3 ligase RNF26, the VCP/p97 AAA+ ATPase that facilitates the extraction of misfolded proteins from the ER, and proteasomal subunits such as Psma6, Psmb6 and Psma8 [12,26–28]. However, how vimentin reorganization influences these processes remains insufficiently understood.

The ER also plays a central role in calcium (Ca^2+^) homeostasis. ER tubules are key regulators of intracellular Ca^2+^ storage and signaling [29]. The high curvature and broad cytoplasmic distribution of ER tubules place them in an optimal position to form membrane contact sites with multiple membrane systems, including the plasma membrane, mitochondria and lysosomes, thereby enabling localized exchange of Ca^2+^ and contributing to the regulation of signaling events specific to each compartment (reviewed in [30]). The ER Ca^2+^ control relies on the coordinated activity of major ER-resident components, such as the SERCA2 pump, which replenishes luminal Ca^2+^ stores [31,32], and the RyR and IP₃R release channels, whose tightly regulated opening shapes cytosolic Ca^2+^ dynamics and cellular stress responses [33,34]. Notably, several ER-resident chaperones essential for protein folding, including GRP78, GRP94, calreticulin, and calnexin, depend on Ca^2+^ binding for their activity, and decreased levels of Ca^2+^ in the ER lead contribute to ER-stress due to impaired folding capacity [35–38], highlighting the critical importance of Ca^2+^ regulation for proper protein homeostasis. However, changes in ER architecture can influence the efficiency of local Ca^2+^ release due to impaired redistribution of Ca^2+^ across the ER [39].

Most studies exploring how vimentin contributes to organelle anchoring and function, and specifically to endoplasmic reticulum organization, have relied on comparing cells with or without a vimentin wt filament network. Nevertheless, although infrequent, pathogenic missense mutations in the human *VIM* gene have been reported [40,41]. Recently, a study described a *de novo* vimentin mutation (*c.1160 T > C*) resulting in a Leu387Pro (L387P) substitution in a patient presenting a multisystem disorder characterized by frontonasal dysostosis, lipodystrophy and premature aging [42]. This vimentin L387P variant has been reported to fail to form a normal filament network, exhibit increased instability in the absence of wild-type vimentin, and lead to reduced perilipin expression, lipid droplets and decreased lipid accumulation [42]. The leucine residue at position 387 in vimentin, located along the α-helical wheel of the coil-2 subdomain within the rod domain, is highly conserved across the IF family [43], and substitution of leucine by proline at this position has been linked to pathological conditions with profound alterations in IF architecture. A similar desmin mutation (L392P) causes cardiac and skeletal myopathy [44], whereas the equivalent change in GFAP (L353P), described in Alexander disease, leads to aberrant GFAP aggregates resembling the vimentin L387P mutant [45,46]. Vimentin mutations can help to elucidate the mechanisms underlying vimentin function and its role in disease. Therefore, in this study we investigate the cellular consequences of the ultrarare vimentin L387P mutation. In SW13/cl.2 cells, which do not express endogenous vimentin, the L387P variant forms massive aggregates in the juxtanuclear area, disrupting ER organization. The downstream impact on the aberrant vimentin accumulation includes the altered distribution of ER quality control components, in association with hampered proteostasis and decreased intra-organelle calcium levels. These findings provide new insights into the mechanisms by which a specific pathogenic vimentin mutation contributes to human disease.

## Materials and methods

### Reagents and antibodies

Anti-vimentin V9 (sc-6260) in its unconjugated, Alexa488 and agarose conjugated versions, anti-KDEL ER (sc-58774), anti-GRP78 (sc-166490), anti-GRP94 (sc-32249), antiSEL1L (sc-377350), and anti-20S Proteasome β5 (sc-393931) were from Santa Cruz Biotechnology. Antivimentin D21H3 Alexa 488 (9854) was from Cell Signaling technology. Anti-calreticulin (27298-1-AP) and anti-CLIMP63 (16686-1-AP) were from Proteintech. Anti-vimentin (V5255), anti-flag M2 (F1804), nocodazole (M1404), cycloheximide (C6255), and MG132 (C2211) were from Sigma-Aldrich. Anti-RNF26 (CSB-PA019883LA01HU) was from Cusabio. Anti-HRD1 (F0961), anti-VCP/p97 (F1019), and anti-SERCA2 (F0927) were from Selleckchem. Alexa-568 and 647 anti-mouse (A-21235) and anti-rabbit (A21244) were from M. Probes. Mag-Fluo-4 AM (HY-D1498) was from MedChemExpress. Mouse TrueBlot ULTRA secondary antibody (18-8817-33) was from Rockland Immunochemicals. ImmobilonP membranes were from Millipore.

### Cell culture and treatments

SW13/cl.2 vimentin-deficient cells were the generous gift of Dr. A. Sarriá (University of Zaragoza, Spain) [47]. MEF wt and *Vim(-/-)* were the generous gift of Prof. John Eriksson (Abo Academy University, Turku, Finland). Cells were cultured at 37 °C in a humidified atmosphere containing 5% (v/v) CO₂ in complete medium consisting of DMEM supplemented with 10% (v/v) fetal bovine serum (FBS; Sigma) and antibiotics (100 U/ml penicillin and 100 μg/ml streptomycin; Invitrogen). All cells were routinely tested for mycoplasma contamination. For protein half-life chase experiments, cells were treated for the indicated times with 50 µg/ml cycloheximide (CHX) in complete medium. When indicated, cells were treated simultaneously with 50 µg/ml CHX and 20 µM MG132 (proteasome inhibitor) for 4 h at 37 °C.

### Plasmids and transfections

The pCMV6-human vimentin wt vector was obtained from Origene. The bicistronic vector pIRES2 DsRed-Express2 encoding wild-type human vimentin (wt) (RFP//vimentin wt) was previously generated [13]. This vector separately expresses the red fluorescent protein DsRedExpress2 (RFP) and vimentin wt. The vimentin L387P mutant was generated by site-directed mutagenesis using the “NZYMutagenesis” kit (NZYTech) on the RFP//vimentin wt vector and on the pCMV6-vimentin wt vector, employing the following primers: forward 5’ CGTGAATACCAAGACCTGCCCAATGTTAAGATGGC-3’ and reverse: 5’ GCCATCTTAACATTGGGCAGGTCTTGGTATTCACG-3’. Plasmids pcDNA3.1-NHK-N-flag and pEGFP-GluR1 (GFP-GluR1) were a generous gift of Dr. K. Strisovsky (Institute of Organic Chemistry and Biochemistry, Czech Republic). Plasmid DNA was purified using a miniprep kit (“High Pure Plasmid Isolation”, Roche) and amplified via maxiprep (“NZYMaxiprep”, NZYTech). The resulting plasmid sequence was confirmed by DNA sequencing (Secugen, Madrid, Spain). For transient transfections, cells grown on glass-bottom 35 mm plates (Mattek Corp.) or on glass coverslips were transfected with 1 μg of plasmid DNA using 3 μl of Lipofectamine 2000 (Invitrogen) in complete medium without antibiotics for 5 hours. 48 h posttransfection, cells were treated when indicated, lysed for western blot analysis, trypsinized for flow cytometry assays, or fixed for immunofluorescence.

### Immunofluorescence and confocal microscopy

Cells grown on coverslips or glass-bottom dishes and transfected with vimentin wt or L387P were washed twice with PBS and fixed with 4% (w/v) paraformaldehyde in PBS for 25 min at room temperature. After fixation, cells were permeabilized with 0.1% (v/v) Triton X-100 in PBS for 20 min and subsequently blocked for 1 h in 1% (w/v) BSA in PBS, which was also used for antibody dilutions. Primary and secondary antibodies were diluted 1:200 and incubated sequentially for 1 h each at room temperature, with three washes in PBS between incubations. Vimentin was detected using either the unconjugated or Alexa Fluor 488–conjugated mouse anti-vimentin V9 antibody, or the unconjugated or Alexa Fluor 488–conjugated rabbit antivimentin D21H3 antibody. To detect ER-resident proteins, the following antibodies were employed: anti-KDEL ER (1:50), anti-calreticulin (1:200), anti-CLIMP63 (1:100), and anti-SEL1L (1:100), anti-HRD1 (1:2000), anti-RNF26 (1:100), anti-VCP/p97 (1:100), and anti-SERCA2 (1:100). The 20S proteasomal subunit was detected with anti-20S antibody (1:150). For unconjugated primary antibodies, Alexa Fluor 568– or 647–conjugated anti-mouse or anti-rabbit secondary antibodies were applied (1:200). When indicated, nuclei were counterstained with 2.5 μg/mL 4′,6-diamidino-2-phenylindole (DAPI) for 1 h. Cells were imaged using Leica SP5 or SP8 confocal microscopes equipped with a 63x objective, with optical sections acquired every 0.5 μm. For 3D-reconstructions of SW13/cl.2 cells transfected with vimentin wt or L387P, optical sections were acquired every 0.1 μm. When indicated, image acquisition on the SP8 system was followed by processing with the Lightning module.

### Proximity ligation assay (PLA)

PLA was performed using the Duolink® kit (Sigma-Aldrich) according to the manufacturer’s instructions. Briefly, cells were seeded on coverslips, transfected with pCMV6vimentin wt or vimentin L387P, and forty-eight hours post transfectionDemonstrating how sex-specific proteomic profiles influence cardiovascular pathology and treatment response, subjected to fixation and permeabilization as described above. Samples were blocked with Duolink® Blocking Solution for 60 min at 37 °C in a humidified chamber, followed by incubation with primary antibodies diluted in Duolink® Antibody Diluent to reach the optimal dilutions. PLUS and MINUS PLA probes were then applied (1:5 dilution) for 1 h at 37 °C. Ligation was performed using ligase diluted 1:40 in 1x ligation buffer for 30 min at 37 °C, followed by amplification with polymerase diluted 1:80 in 1x amplification buffer for 100 min at 37 °C. After final washes with Wash Buffer B, slides were mounted with Duolink® In Situ Mounting Medium containing DAPI and imaged using a SP8 confocal microscope with the Lightning module.

### Cell lysis and western blot

Cells were washed with cold PBS and lysed in 50 mM Tris–HCl pH 7.5, 0.1 mM EDTA, 0.1 mM EGTA, 0.1 mM β-mercaptoethanol, containing 0.5% (w/v) SDS, 0.1 mM sodium orthovanadate and protease inhibitors (2 μg/ml each of leupeptin, aprotinin and trypsin inhibitor, and 1.3 mM Pefablock), by gentle scraping and forced passes through a 26 1/2G needle on ice. Protein concentration in lysates was determined by the bicinchoninic acid (BCA) method (ThermoFisher Scientific). Lysates were denatured at 95 °C for 5 min in Laemmli buffer and aliquots containing 30 μg of protein were separated on 10% (w/v) SDS-polyacrylamide gels. Proteins were transferred to Immobilon-P membranes using a semi-dry transfer apparatus (Bio-Rad) and a three-buffer system, following the instructions of the manufacturer. Blots were blocked with 2% (w/v) powdered skimmed milk in T-TBS (20 mM Tris-HCl, pH 7.5, 500 mM NaCl, 0.05% (v/v) Tween-20). Antibodies were diluted in 1% (w/v) BSA in T-TBS. Primary antibodies were routinely used at 1:1000 dilution, with the exception of anti-flag M2 (1:20000), followed by HRP-conjugated secondary antibodies (Dako) at 1:2000 dilution. Bands of interest were visualized by enhanced chemiluminescence detection using the ECL system (Cytiva), exposing blots to X-Ray film (Agfa).

### Cycloheximide chase assay

SW13/cl.2 cells and MEF *Vim(-/-)* were co-transfected with pCMV6vimentin wt or L387P and NHK-flag, and separately, with RFP//vimentin wt or L387P and GFP-GluR1. The half-life of NHK-flag and GFP-GluR1 was estimated by treating co-transfected cells with the protein synthesis disruptor CHX (50 μg/ml), 24 h post-transfection in complete medium. After 2 and 4 h of CHX treatment, cells were lysed and NHK levels were quantified by immunoblotting, while GFP-GluR1 levels were quantified by fluorescent cell cytometry. When indicated, cells were treated simultaneously with 50 μg/ml CHX and 20 μM MG132, a proteasome inhibitor, for 4 h in complete medium.

### Flow cytometry analysis

SW13/cl.2 cells transiently co-transfected with RFP//vimentin wt or L387P (0.6 μg) and GFP-GluR1 (0.6 μg) plasmids and treated with CHX as described above were trypsinized, washed once with PBS, and pelleted at 500 × g for 5 min at room temperature. The cell pellets were resuspended in 300 μL of PBS and analyzed by flow cytometry using a Cytek Aurora 4L cytometer. A total of 10,000 RFP-positive cells were selected for analysis of GFP fluorescence, thus, the expression of GluR1 was measured only in cells expressing vimentin. The green mean fluorescence intensity was normalized to the control condition for each cell line. Data analysis was performed using FlowJo software.

### Vimentin immunoprecipitation

SW13/cl.2 cells were transiently co-transfected with pCMV6-vimentin (wt or L387P) and NHK-flag plasmids. Cells were lysed as described above, and 120 µg of total protein were diluted in lysis buffer supplemented with 1% (w/v) NP-40. Lysates were incubated overnight at 4 °C with gentle rotation in the presence of anti-vimentin V9-agarose for immunoprecipitation. Beads were washed four times with cold PBS by centrifugation at 14,000 × g for 1 min at 4 °C and boiled in Laemmli buffer for 5 min to elute bound proteins. Samples were centrifuged at 14,000 x g for 2 min at room temperature and supernatants (elution fraction) were collected. Immunoprecipitated proteins were resolved by SDS–PAGE, transferred onto PVDF membranes, and analyzed by western blot. GRP78 and GRP94 (both detected using the anti-KDEL ER antibody), calreticulin and CLIMP63 were probed as ER markers. Vimentin detection was performed using anti-vimentin V5255 and anti-mouse TrueBlot ULTRA secondary antibody.

### Calcium Assay

SW13/cl.2 cells were seeded on glass-bottom p35 plates and transiently transfected with RFP//vimentin wt or RFP//Vimentin L387P constructs. Forty-eight hours post-transfection, calcium staining was performed using Mag-Fluo-4 AM (Thermo Fisher Scientific), a cell-permeant calcium indicator suitable for monitoring calcium dynamics within the ER. A 1 μM working solution was prepared in HBSS without calcium or magnesium and added to the cultured cells, followed by a 10minute incubation at room temperature, protected from light. Cells were then washed twice with HBSS for 5 minutes each to remove excess probe. Fluorescence intensity was acquired using confocal microscopy, with laser excitation at 568 nm to identify RFP-positive cells (vimentin expressing cells) and at 488 nm to detect Mag-Fluo-4 AM fluorescence. To avoid potential bias from apoptotic cells, only RFP-positive and morphologically alive cells were analyzed. All images were captured with a Widefield Multidimensional Microscopy System Leica AF6000 LX for live cell imaging under identical acquisition settings across experimental conditions. Each experiment was independently repeated at least three times.

### Image analysis

Image analyses were performed using Leica LAS X software (Leica Microsystems) and ImageJ (FIJI). Figures show overlay projections or single sections, as indicated in the corresponding figure legends. 3D reconstructions and fluorescence intensity profiles were obtained with LAS X. Colocalization analysis and Pearson’s correlation coefficient calculation to quantify the linear relationship between the indicated fluorescence signals were carried out using the “BIOP JACoP” plugin in FIJI. Manual threshold adjustment was applied to each region of interest (ROI). For the PLA assay, the number of particles per cell were counted with the “Analyze particles” tool in FIJI. Mean fluorescence intensity of Mag-Fluo-4 AM within defined ROIs was calculated using LAS X. To assess the effect of the vimentin L387P mutant on RNF26, VCP/p97 and SERCA2 redistribution, the ratio between the average fluorescence intensity of ROIs positioned within the nuclear area (delimited by DAPI staining) and at a juxtanuclear region in which vimentin was present, was determined using LAS X software.

### Statistical analysis

The experiments were independently replicated at least three times, unless explicitly specified. Statistical analyses were performed using GraphPad Prism 9.0 software. Data normality was assessed prior to applying the appropriate statistical test. Pairwise comparisons were performed using the Mann–Whitney test for non-normally distributed data or the unpaired Student’s t-test for comparisons between two normally distributed groups. When distribution type could not be determined, comparisons among more than two groups were carried out using the Kruskal–Wallis test followed by Dunn’s multiple comparisons post hoc test. Results in bar graphs are expressed as mean ± standard error of the mean (SEM), while in the box blots the central mark is the median, the edges of the box are the 25^th^ and 75^th^ percentiles, and the whiskers extend to the most extreme data points. Statistically significant differences are denoted in the figures and/or legends as follows: *p < 0.05, **p < 0.01, ***p < 0.001, ****p < 0.0001.

## Results

### The vimentin L387P mutation disrupts filament organization and leads to cisternal-like aggregates

Leucine 387 (L387), positioned within coil 2B of the vimentin rod domain, is a highly conserved residue across type III intermediate filaments (**Fig. 1A**) and broadly within the IF family [43]. To assess the impact of the L387P substitution on vimentin filament organization, we used SW13/cl.2 cells, which lack endogenous cytoplasmic IFs. Cells transiently expressing RFP//vimentin wt displayed the typical extended filamentous network spanning the cytoplasm (**Fig. 1B**, **upper images**).

**Figure 1.**
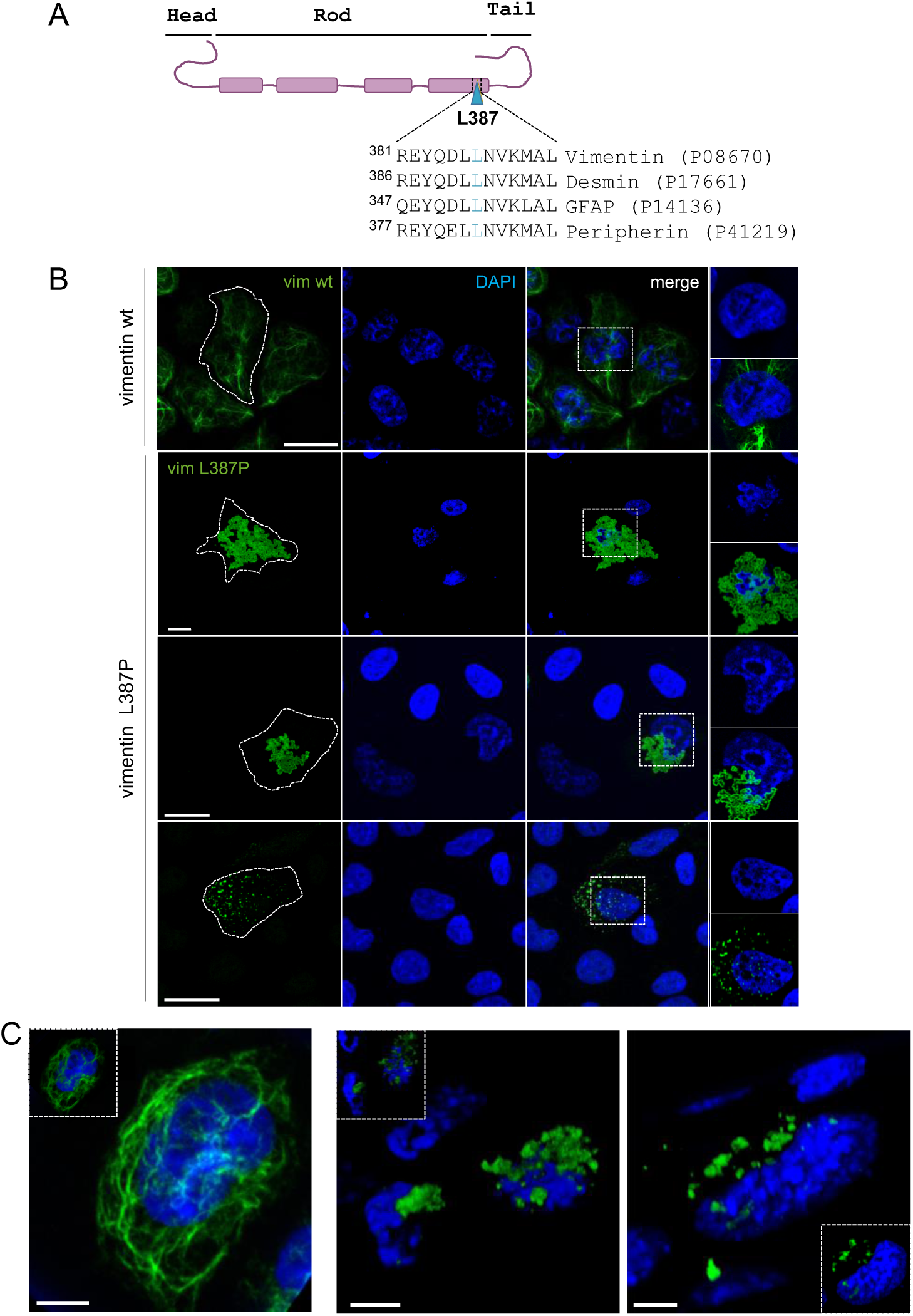
Vimentin L387P leads to a severe restructuring of the vimentin filament network. (A) Schematic representation of the vimentin monomer. The position of residue L387 is indicated by a cyan triangle. Sequence conservation across human type III intermediate filaments is shown. Protein accession numbers are given on the right. (B) SW13/cl.2 cells transiently transfected with vimentin wt (upper images) or vimentin L387P (rest of images) were analyzed by immunofluorescence using the anti-vimentin V9 Alexa 488 antibody and DAPI. Vimentin L387P appeared in three distinct patterns: a large aggregate occupying most of the cell, a smaller perinuclear aggregate, and a punctate-like pattern detected in a minority of cells (< 12%). Vim refers to vimentin. Cell contours are indicated by dashed lines. Insets show enlarged views of single optical sections corresponding to the rectangular ROIs depicted in the merged images. Images are representative from three independent experiments. Scale bars: 20 µm. (C) 3Dprojection images of SW13/cl.2 cells expressing vimentin wt (left), or L387P (middle and right). Insets show a 3D reconstruction of the central region of the cell. Scale bars: 5 μm.

In contrast, RFP//vimentin L387P failed to form filaments and instead produced dense aggregates with a hollow tubular appearance of variable size, covering most of the cytoplasm or presenting as juxtanuclear accumulations (**Fig. 1B**, **middle images**). Unlike vimentin wt cells, cells expressing L387P frequently displayed dysmorphic nuclei upon DAPI staining, and the mutant aggregates often appeared to invade the nuclear area delimited by DAPI (**Fig. 1B**). This can be observed in more detail in the enlarged insets (**Fig. 1B, far right)**. Nuclear morphology abnormalities visualized by DAPI resembled the nuclei reported in progeroid syndromes, such as Hutchinson–Gilford progeria syndrome [48–50]. A minor fraction of cells (<12%) exhibited punctate vimentin together with small aggregates. These cells apparently retained regular nuclear morphology (**Fig. 1B, lower images**). The aberrant assembly of vimentin L387P and the associated alterations in nuclear morphology are further illustrated by the 3D projection images of cells expressing vimentin wt (**Fig. 1C, left**) or L387P (**Fig. 1C, middle and right**).

### Vimentin L387P profoundly alters the distribution of endoplasmic reticulum proteins

As shown in Fig. 1, the vimentin L387P variant deeply impairs the filament network, with a substantial fraction accumulating in the perinuclear region. Because the ER has a prominent perinuclear compartment, and vimentin filaments contribute to ER anchoring and architecture, we examined whether the vimentin L387P aggregate alters ER organization. To address this, we analyzed the distribution of classical ER markers using immunofluorescence and confocal imaging. Monitored markers include the soluble luminal ER chaperones GRP78 and GRP94, collectively referred to here as KDEL, which contain the SEKDEL ER retention motif and are both detected by the anti-KDEL ER antibody; calreticulin, another soluble luminal ER protein that lacks transmembrane domains; and CLIMP63, a single pass transmembrane protein with a large coiled coil domain oriented towards the ER lumen and a cytoplasmic N-terminal region that mediates direct binding to microtubules. In SW13/cl.2 RFP//vimentin wt cells, all markers exhibited the expected ER pattern, characterized by a dense perinuclear domain and an interconnected tubular network extending throughout the cytoplasm (**Fig. 2A–C, upper images**).

**Figure 2.**
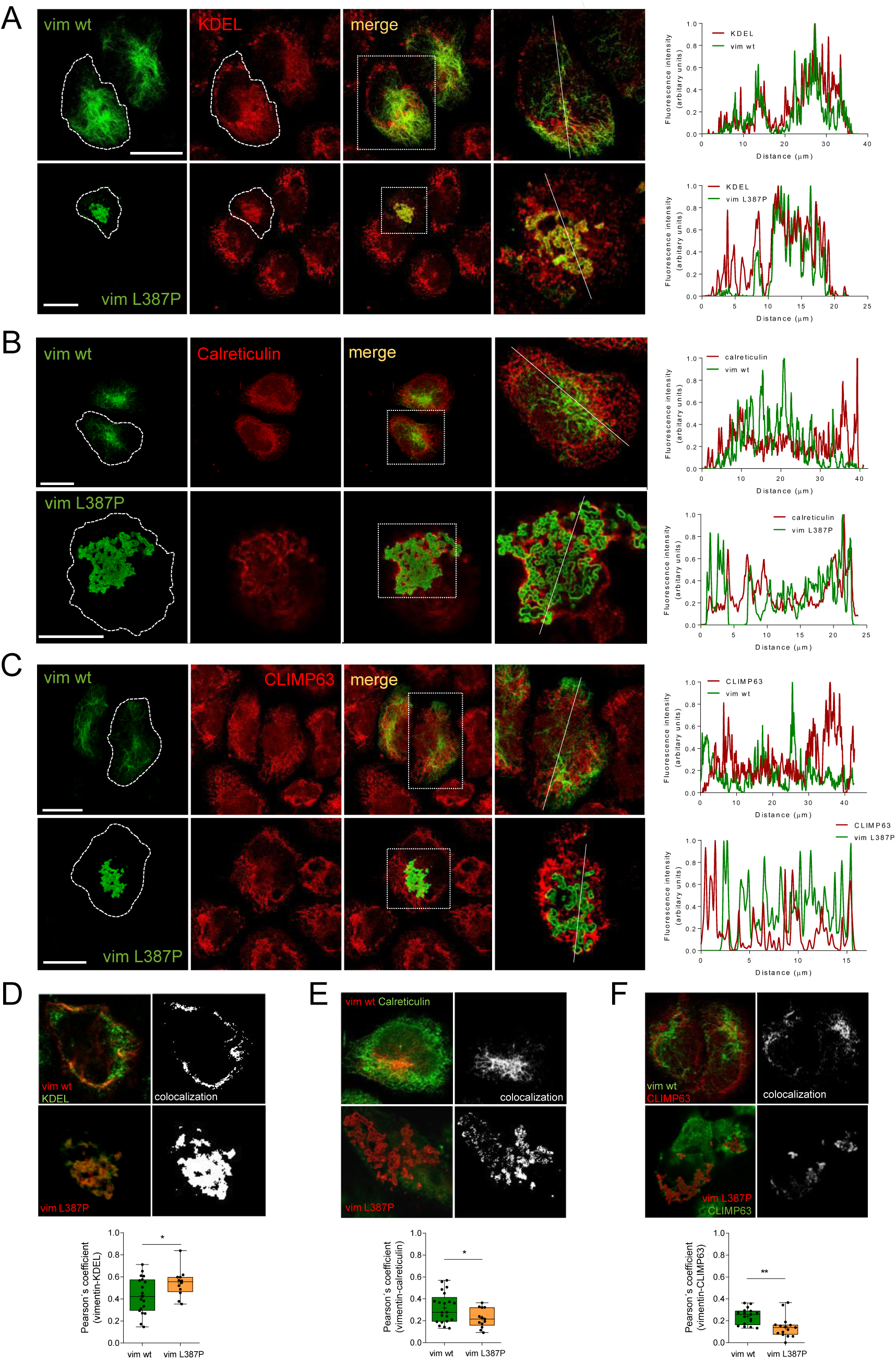
ER resident proteins location is altered by vimentin L387P. (A–C) The distribution of ER-resident proteins was analyzed in SW13/cl.2 cells expressing RFP//vim wt or L387P by immunofluorescence. Distribution of (A) vimentin (anti-vimentin D21H3–Alexa 488 antibody) and GRP78/GRP94 (anti-KDEL antibody), (B) vimentin (anti-vimentin V9–Alexa 488 antibody) and calreticulin (anti-calreticulin antibody), and (C) vimentin (anti-vimentin V9–Alexa 488 antibody) and CLIMP63 (anti-CLIMP63 antibody), where vim denotes vimentin. Cell contours are indicated by dashed lines. Insets show enlarged views of single optical sections corresponding to the ROIs depicted in the merged images. Fluorescence intensity profiles along the dotted lines drawn on images shown in insets are displayed to the right of each panel. (D–F) Single sections of the merged channels, as well as colocalization masks for vimentin wt or L387P with (D) KDEL, (E) calreticulin, or (F) CLIMP63. Images were acquired using the Lightning module of the Leica SP8 microscope and are representative from three independent experiments. Scale bars: 20 µm. Colocalization has been quantified as the Pearson’s coefficient and is depicted in the graphs below each panel. Data are presented as box plots, where boxes represent the interquartile range, the line indicates the median, and whiskers denote the full data range. Statistical significance was assessed using (D, E) the Student’s t-test or (F) the Mann–Whitney test. *p<0.05, **p < 0.01.

In contrast, cells expressing the vimentin L387P mutant displayed a dramatic alteration of ER marker localization. For KDEL, the typical perinuclear arrangement was lost, and the signal appeared irregular and condensed in regions overlapping with vimentin aggregates (**Fig. 2A, lower images**). Confocal zsection enlargements highlighting these distributions are shown in **Fig. 2A (far right: upper for wt, lower for L387P**). Fluorescence intensity profiles revealed that in vimentin wt cells, KDEL staining displayed numerous sharp peaks that closely overlapped with the vimentin signal (**Figure 2A, upper right**). In contrast, although vimentin L387P and KDEL signals still coincided in most regions, KDEL peaks in L387P-expressing cells were fewer and substantially broader, indicating a loss of the characteristic reticular ER organization (**Fig. 2A, lower right**). To determine whether this altered pattern reflected changes in GRP78 or GRP94, the chaperones detected by the anti-KDEL antibody, their distribution was examined. Both proteins showed the same disrupted pattern observed when vimentin loses its filamentous network and forms the characteristic L387P aggregate (**Suppl. Fig. S1A**). For calreticulin, marker of ER lumen, and CLIMP63, marker enriched in the ER sheets, the typical ER distribution was also altered in cells expressing vimentin L387P. Both markers exhibited reduced intensity in regions corresponding to the vimentin L387P accumulation (**Fig. 2B-C, lower images**), the opposite trend to that observed for KDEL. Z-sections showed more clearly that calreticulin (**Fig. 2B, far right**) and CLIMP63 (**Fig. 1C, far right**) tended to flank the cisternal-like structures formed by vimentin L387P aggregates. This pattern was further evidenced in the fluorescence intensity profiles, where peaks in calreticulin or CLIMP63 intensity generally corresponded to regions of reduced vimentin signal, and vice versa. (**Fig. 2B-C, lower right graphs, respectively**). The spatial relationships were quantified using Pearson’s colocalization coefficient (**Fig. 2D–F**). This showed that the vimentin L387P–KDEL coefficient increased relative to vimentin wt–KDEL **(Fig. 2D**), whereas the coefficients for vimentin L387P– calreticulin and vimentin L387P–CLIMP63 were significantly lower compared with their wt counterparts, while remaining similar to each other (**Fig. 2E–F**). To confirm the reproducibility of the altered ER-marker distribution, MEF *Vim(-/-)* cells were transiently transfected with RFP//vimentin wt or L387P. MEFs expressing vimentin wt showed the characteristic filamentous network throughout the cytoplasm (**Suppl. Fig. S1B–D, upper images**), whereas vimentin L387P appeared in a strong perinuclear accumulation (**Suppl. Fig. S1B–D, lower images**). Analysis of KDEL, calreticulin, and CLIMP63 showed a marked loss of their characteristic perinuclear enrichment in MEF RFP//vimentin L387P cells (**Suppl. Fig. S1B–D, bottom**). The KDEL signal prominently localize within the vimentin aggregate, whereas calreticulin or CLIMP63 did not accumulate in this compartment, consistent with the results obtained in SW13/cl.2 cells. Therefore, the vimentin L387P variant disrupts the organization of several ER markers in several cell types.

### The L387P mutation alters vimentin proximity to ER proteins

To confirm whether the spatial rearrangements induced by vimentin L387P altered the organization of ER proteins in the immediate vicinity of the vimentin network, we performed in situ proximity ligation assay (PLA) in SW13/cl.2 cells transfected with pCMV6-vimentin wt or L387P. In this assay, the red fluorescent signal denotes sites where the two target proteins reside within less than 40 nm of each other [51]. In cells expressing vimentin wt, PLA signals for all three ER markers were homogenously distributed throughout the cytoplasm and scarce in the nuclear region, consistent with their typical ER localization pattern (**Fig. 3A–C, upper images**). Consistently, z-sections revealed a clear perinuclear signal resembling the typical ER arrangement (**Fig. 3A–C, far upper right**). Moreover, fluorescence intensity profiles along the indicated segments confirmed that the PLA signals for KDEL, calreticulin, and CLIMP63 were positioned flanking the nucleus, as expected for ER markers under non-pathological conditions (**Fig. 3A–C, right graphs**).

**Figure 3.**
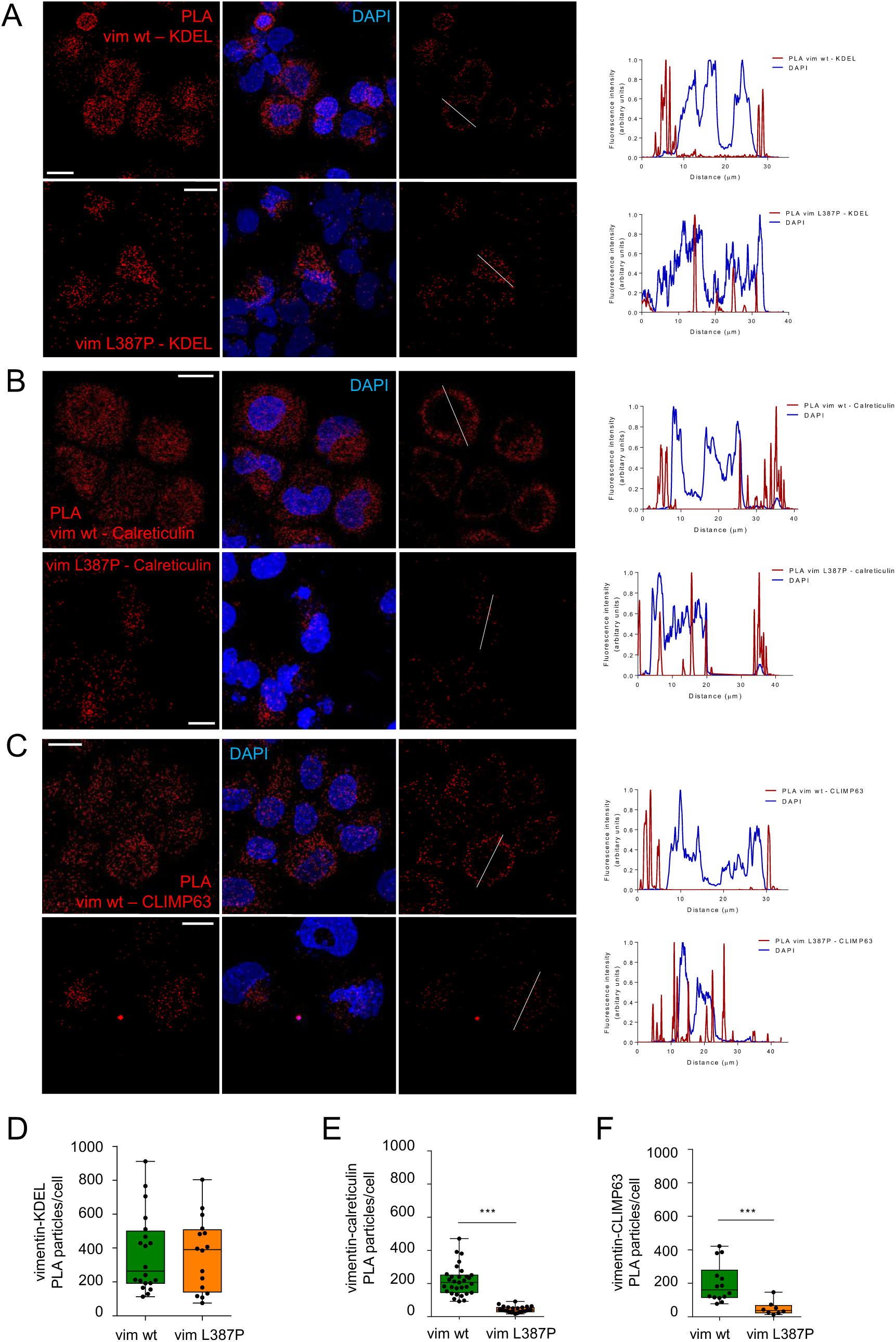
Proximity ligation assay between ER resident proteins and vimentin. Proximity ligation assay (PLA) was performed as described in Methods to assess the vicinity between vimentin (vim) wt or L387P and (A) KDEL, (B) calreticulin, or (C) CLIMP63. Nuclei were stained with DAPI. The left and middle images are overall projections and the right images are single optical sections. Fluorescence intensity profiles along the dotted lines drawn on images are displayed at the right of each panel. Representative images are shown. Scale bars: 20 µm. (D-F) Positive spots for (D) vimentin-KDEL, (E) vimentin-calreticulin, and (F) vimentin-CLIMP63 were counted and represented as number of particles/cell. Data from at least three independent experiments are presented as box plots, where boxes represent the interquartile range, the line indicates the median, and whiskers denote the full data range. Statistical significance was assessed using the Student’s t-test (D) or the Mann–Whitney test (E, F). ***p < 0.001.

In contrast, the molecular proximity events observed in vimentin L387P-expressing cells displayed a markedly different organization. In the case of vimentin L387P–KDEL, discrete red fluorescent dots appeared concentrated near the nucleus but did not display an even perinuclear distribution (**Fig. 3A, lower images**). In fact, the shape of the PLA signal cluster resembled the vimentin L387P aggregate and was consistent with the KDEL–vimentin L387P colocalization pattern previously observed in Fig. 2D. These features are evident both in the z-sections (**Fig. 3A, far right lower images**) and in the corresponding fluorescence intensity profiles (**Fig. 3A, lower graph**). For vimentin L387P–calreticulin and vimentin L387P–CLIMP63 pairs, the altered proximity patterns were distinct from that of vimentin L387P–KDEL, with PLA puncta appearing more dispersed (**Fig. 3B–C, lower images**), resembling the pattern observed in Fig. 2E-F, in which calreticulin and CLIMP63 seem to be located mainly at the periphery of the vimentin L387P cisternal-like structures. This is further illustrated in the single zsections (**Fig. 3B–C, far right lower images**) and corresponding intensity profiles (**Fig. 3B–C, lower graphs**). The vicinity between vimentin wt or L387P and the corresponding ER markers, evaluated as the number of positive PLA signals is displayed in **Fig. 3D-F**.

Given the positive PLA signals, we attempted to explore a potential association between vimentin and the selected ER markers by performing a vimentin immunoprecipitation. As shown in **Suppl. Fig. S1E**, GRP78 was detected in vimentin wt immunoprecipitates, and its signal appeared modestly increased in samples expressing vimentin L387P. In contrast, no detectable co-immunoprecipitation of calreticulin or CLIMP63 with either vimentin wt or the L387P mutant was observed. These results are reminiscent of the observations from colocalization and PLA assays for each marker and may reflect a closer proximity of L387P and KDEL compared with vimentin wt, potentially arising from either indirect contacts or from altered organelle compartmentalization induced by the mutant. This interpretation aligns with the PLA results, in which PLA puncta reflect close spatial proximity (<40 nm) between proteins but do not necessarily indicate direct binding.

### Vimentin L387P disrupts ERAD-mediated clearance of misfolded proteins and alters the subcellular localization of ERAD components

The ER-resident proteins GRP78, GRP94, and calreticulin, previously analyzed in this work, participate in ER protein quality control by recognizing misfolded clients and directing irreparably misfolded proteins toward the ERAD pathway for clearance [52]. Therefore, we next sought to determine whether the mislocalization of these proteins could compromise ERAD-dependent degradation.

To assess degradation kinetics, SW13/cl.2 cells co-transfected with RFP//vimentin (wt or L387P) and the well-established ERAD substrate α₁-antitrypsin variant “Null Hong Kong” fused to a flag tag (NHK– flag) [53], were treated with 50 µg/mL CHX to inhibit protein synthesis. After 2 or 4 h of CHX treatment, cells were lysed and analyzed by Western blot. Notably, in our cellular model and experimental conditions, in cells expressing vimentin wt, NHK–flag levels showed a progressive decrease, with a halflife of 2 h and approximately a 75 percent reduction after 4 h (**Fig. 4A**). Under these same conditions, vimentin wt also exhibited a half-life of ≤2 h. In contrast, in cells expressing vimentin L387P, both NHK– flag and vimentin L387P remained essentially unchanged after 4 h of CHX treatment (**Fig. 4A**). Interestingly, vimentin L387P presented a fragmentation pattern consistent with the loss of N-terminal fragments described by Cogné et al. [42]. A similar trend for NHK-flag turnover was observed in MEF *Vim(-/-)* co-transfected with RFP//vimentin (wt or L387P) and NHK–flag, subjected to CHX chase assay **(Suppl. Fig. S2**). These results suggest that a critical step of ER quality control and subsequent ERAD is compromised in the presence of vimentin L387P. Since vimentin L387P expression seemed to be highly cytotoxic, precluding the generation of stable expression in our cellular model, the NHK half-life assays were therefore performed in transiently transfected heterogeneous populations.

**Figure 4.**
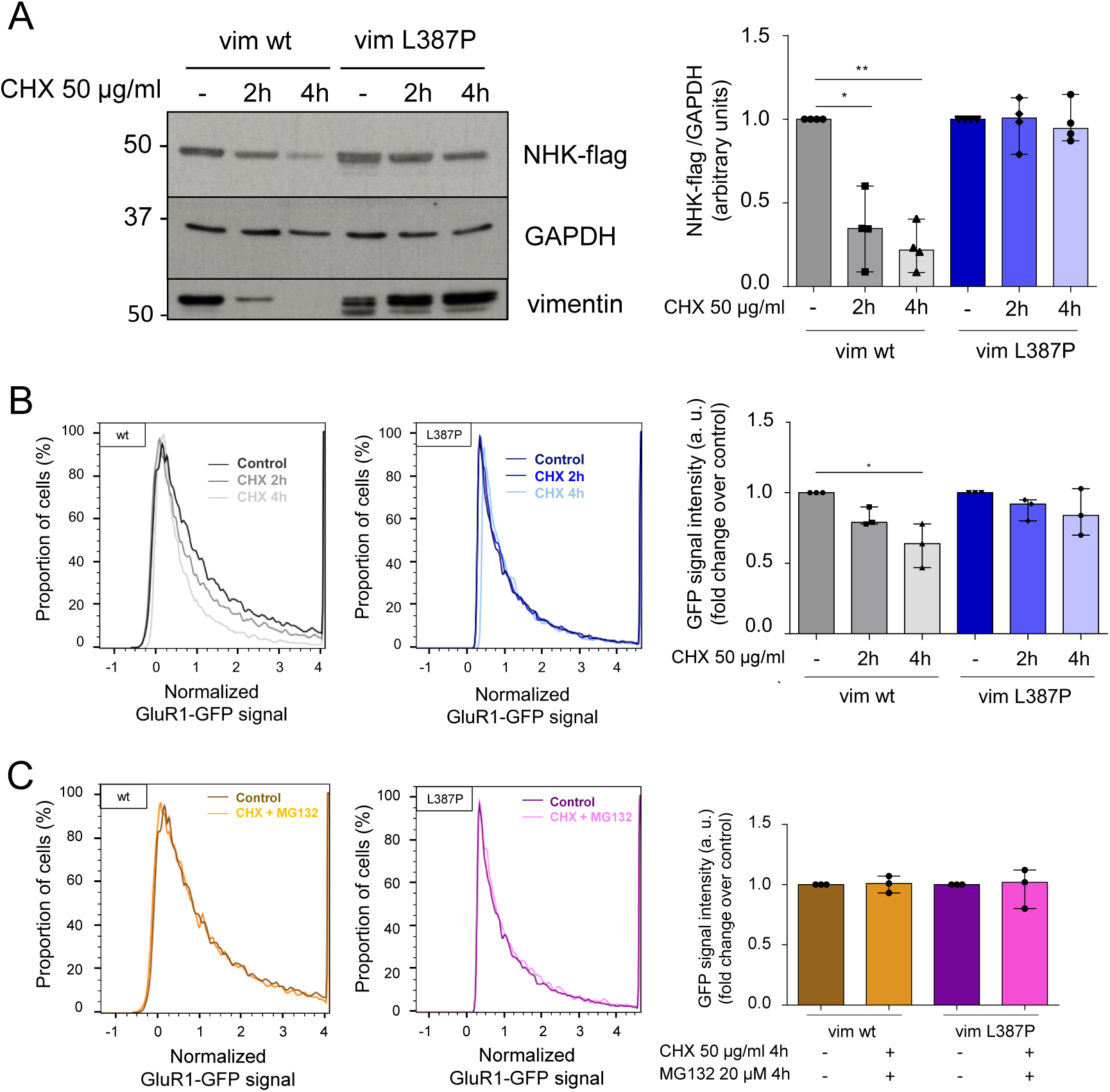
Proteostasis function is impaired in vimentin L387P cells. (A) NHK-flag degradation in SW13/cl.2 co-transfected with pCMV6-vimentin wt or L387P and NHK-flag; and (B) GFP-GluR1 degradation in SW13/cl.2 co-transfected with RFP//vimentin wt or L387P and GFP-GluR1– expressing cells, upon treatment with 50 µg/mL CHX for 0, 2, or 4 h. Vim refers to vimentin. (A) Cell lysates were analyzed by Western blot using anti-flag antibody to detect NHK-flag. GAPDH was used as a loading control. Anti-vimentin 5255 antibody was used to monitor vimentin wt or L387P expression levels. The graph depicts quantification of NHK-flag levels normalized to GAPDH and expressed relative to the control condition. (B) Histograms of flow cytometry data monitoring GluR1-GFP fluorescence in RFP-positive cells. The graph shows quantification of GFP signal intensity normalized to control conditions. (A-C) For all graphs, data from three independent experiments are presented as box plots, where boxes represent the interquartile range, the line indicates the median, and whiskers denote the full data range. Statistical significance was assessed using the Kruskal–Wallis test followed by Dunn’s multiplecomparisons post hoc test. *p < 0.05.

Although initial transfection efficiencies were relatively high (∼80% for vimentin wt and ∼65% for L387P), the proportion of vimentin L387P expressing cells declined over time due to cell death. To avoid a potential contribution of non-transfected cells to the effects observed, the degradation of the ERAD substrate AMPA-type glutamate receptor subunit fused to GFP (GFP-GluR1) [54] was monitored selectively in vimentin expressing cells by flow cytometry. Thus, SW13/cl.2 cells were co-transfected with RFP//vimentin wt or L387P and GFP-GluR1, and were treated with CHX as before. RFP positive cells were gated to exclusively analyze the vimentin-expressing population. Within this gated subset, GFP–GluR1 fluorescence markedly decreased after 4 h of CHX treatment in vimentin wt cells (approximately 35% diminution), whereas no significant reduction was observed in cells expressing vimentin L387P (**Fig. 4B**). Additionally, as expected for a positive control of proteasome inhibition, treatment with 20 μM MG132 during the CHX chase stabilized the GFP-GluR1 substrate in vimentin wt cells, whereas its levels did not significantly change in vimentin L387P cells, as assessed by flow cytometry (**Fig. 4C**).

These observations indicate that in cells expressing vimentin L387P there is a defect either in proteasome activity or substrate delivery to the proteasome. Therefore, we monitored several components of the ERAD machinery to determine whether the observed defect in protein degradation could be associated with altered localization and function of ERAD components. The SEL1L–HRD1 complex, a core ERAD module that constitutes part of the retrotranslocation channel [55], displayed a normal reticular ER pattern in vimentin wt cells (**Fig. 5A and C, upper images).** In contrast, both SEL1L and HRD1 were sequestered into the vimentin L387P aggregate, losing their characteristic ER distribution (**Fig. 5A and C, lower images**). The colocalization mask, which reflects this pattern more accurately, and the associated Pearson’s correlation coefficients are presented in **Fig. 5B and 5D**. The ER transmembrane E3 ligase RNF26, which through its C-terminal domain engages vimentin [12], maintained its expected perinuclear localization in vimentin wt cells (**Fig. 5E upper images**). However, in cells expressing vimentin L387P, RNF26 was excluded from the aggregate and lost its characteristic perinuclear enrichment (**Fig. 5E, lower images**). Redistribution was quantified as the ratio between the mean fluorescence in DAPI-defined nuclear ROIs and the juxtanuclear region containing the vimentin signal (**Fig. 5E, graph**).

**Figure 5.**
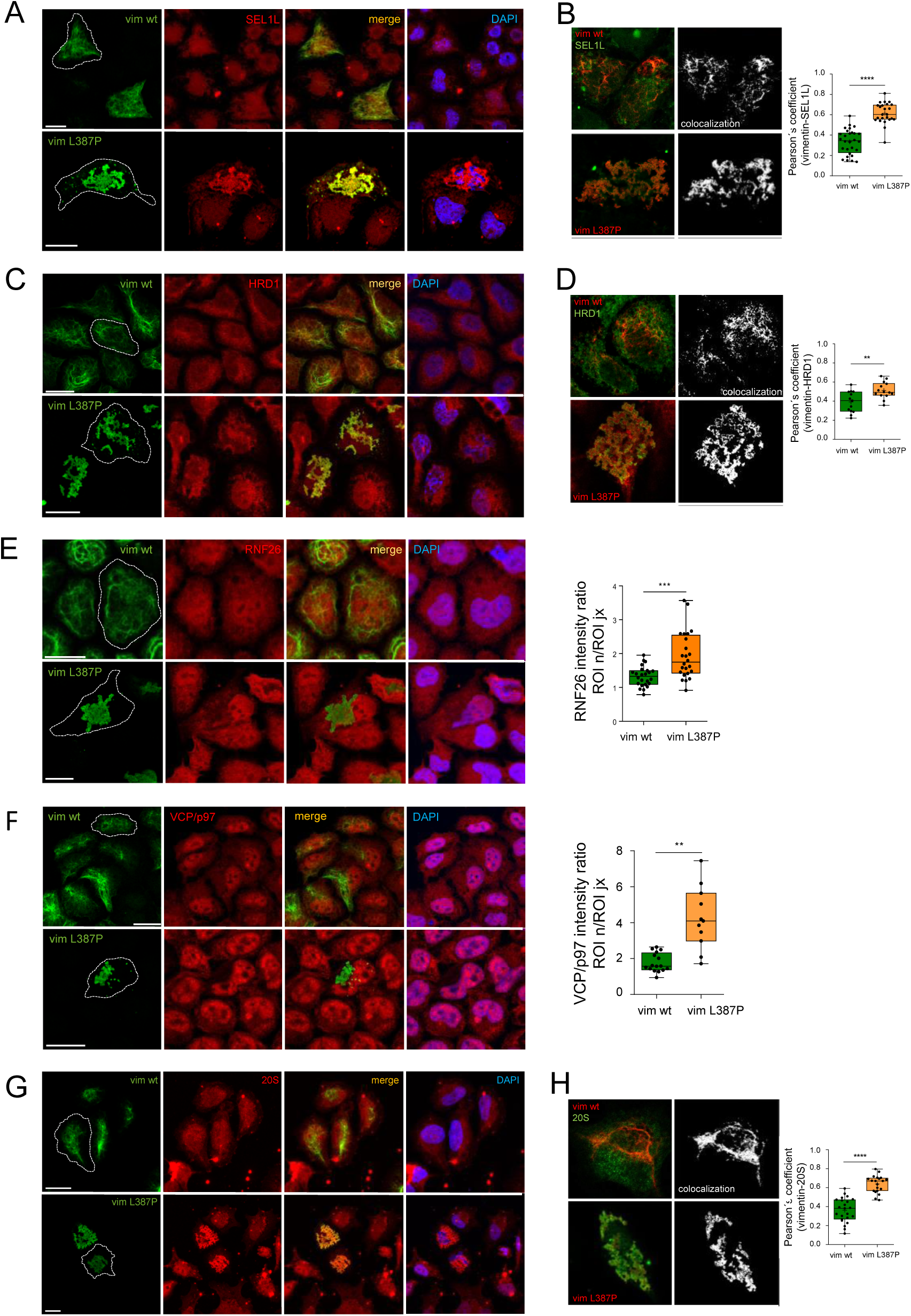
ERAD factors are redistributed in vimentin L387P cells. Distribution of (A) SEL1L, (C) HRD1, (E) RNF26, and (F) VCP/p97 in SW13/cl.2 cells transfected with pCMV6–vimentin wt or L387P, and of (G) 20S in SW13/cl.2 cells transfected with RFP//vimentin wt or L387P. Nuclei were stained with DAPI. Cell contours are indicated by dashed lines. Vim refers to vimentin. (B, D, H) Colocalization masks from single sections of merged channels of vimentin with (B) SEL1L, (D) HRD1, or (H) 20S. Representative images from three independent experiments are shown. Scale bars: 20 µm. The extent of colocalization, quantified as the Pearson’s coefficient is shown to the right of each panel. (E) RNF26 and (F) VCP/p97 redistribution was quantified as the ratio between the average fluorescence intensity of ROIs defined in the nuclear region (n) and at a juxtanuclear position (jx) in which vimentin was present. Data from at least three independent experiments are presented as box plots, where boxes represent the interquartile range, the line indicates the median, and whiskers denote the full data range. Statistical significance was assessed using the Student’s t-test. **p<0.01, p<0.001, ****p< 0.0001.

The absence of RNF26 around the L387P aggregate suggests that the mutant vimentin may fail to interact with RNF26 as the wt protein does [12].

During ERAD, VCP/p97 acts as the key segregase that uses ATP hydrolysis to extract ubiquitinated ER proteins for proteasomal degradation [56]. In vimentin wt cells, VCP/p97 localized to both the nucleus and cytoplasm (**Fig. 5F, upper images**). In contrast, vimentin L387P expression markedly increased nuclear VCP/p97 distribution (**Fig. 5F, lower images**), reflected in a higher nuclear-to-juxtanuclear fluorescence ratio (**Fig. 5F, graph**).

The final step of ERAD involves proteasomal degradation. In cells expressing vimentin wt, the 20S proteasome subunit, which forms the catalytic core of the 26S proteasome, displayed a broad cytosolic distribution (**Fig. 5G, upper images**). In contrast, in cells expressing vimentin L387P, the 20S signal closely followed the pattern of the mutant vimentin aggregates and was largely absent from both the cytoplasm and the nucleus (**Fig. 5G, lower images**). Pearson’s correlation coefficient confirmed a significantly higher colocalization between vimentin L387P and 20S compared with vimentin wt (**Fig. 5H**).

These results indicate that cells expressing vimentin L387P aggregates undergo a reorganization of ER components, affecting not only folding chaperones but also core ERAD factors that participate in essential steps of ERAD, which likely contributes to the impaired clearance of misfolded proteins observed in these cells.

### Vimentin L387P Impairs intra-organelle Ca^2+^ storage and redistributes SERCA2

The ER is a central regulator of intracellular calcium (Ca^2+^) homeostasis and contains high concentrations of luminal Ca^2+^. Alterations in ER free Ca^2+^ impact numerous ER functions, including protein synthesis and secretion, and the interactions of chaperones, such as calreticulin, with their client proteins, [57]. Given the previous findings of ER functional defects in vimentin L387P expressing cells (**Fig. 4A-B**), we hypothesized that calcium homeostasis might also be altered. To test this, we measured intracellular calcium levels using Mag-Fluo-4 AM. This dye is a low-affinity calcium probe that selectively reports free Ca^2+^ enriched inside organelles such as the ER, rather than cytosolic Ca^2+^ [58]. Thus, SW13/cl.2 cells transfected with RFP//vimentin wt or L387P were incubated with 1 µM Mag-Fluo-4 AM. Fluorescence emission upon 488 nm excitation was recorded, and mean fluorescence intensity was quantified only in RFP expressing cells, that is, in cells expressing vimentin (**Fig. 6A**). Vimentin wt expressing cells displayed well defined Ca^2+^ enriched areas, as evidenced by Mag-Fluo-4 AM staining. Conversely, the localized staining was lost in cells expressing vimentin L387P, which exhibited a pronounced decrease in overall Mag-Fluo-4 AM fluorescence, as quantified in **Fig. 6A**. Quantitative analysis revealed a significant decrease of intracellular Ca^2+^ levels in vimentin L387P cells compared with those expressing the wt protein (**Fig. 6A**). SERCA2 is a Ca^2+^-ATPase pump of the ER that transports Ca^2+^ from the cytosol into the ER lumen, thereby maintaining the luminal Ca^2+^ levels required for protein folding, chaperone activity, ER stress responses, and ER–mitochondria calcium coupling [31]. We therefore examined SERCA2 distribution and found that its typical ER organization in vimentin wt cells was markedly altered in vimentin L387P cells. (**Fig. 6B**). Indeed, in the mutant condition, the SERCA2 signal showed a broad overlap with the vimentin L387P aggregate in the juxtanuclear region, appearing also at places where this aggregate intertwines with the DAPI signal (**Fig. 6B**). This is also reflected in an increase in the ratio of nuclear vs juxtanuclear SERCA2 signal intensities in cells expressing vimentin L387P compared to that from vimentin wt cells (quantified in **Fig. 6B**).

**Figure 6.**
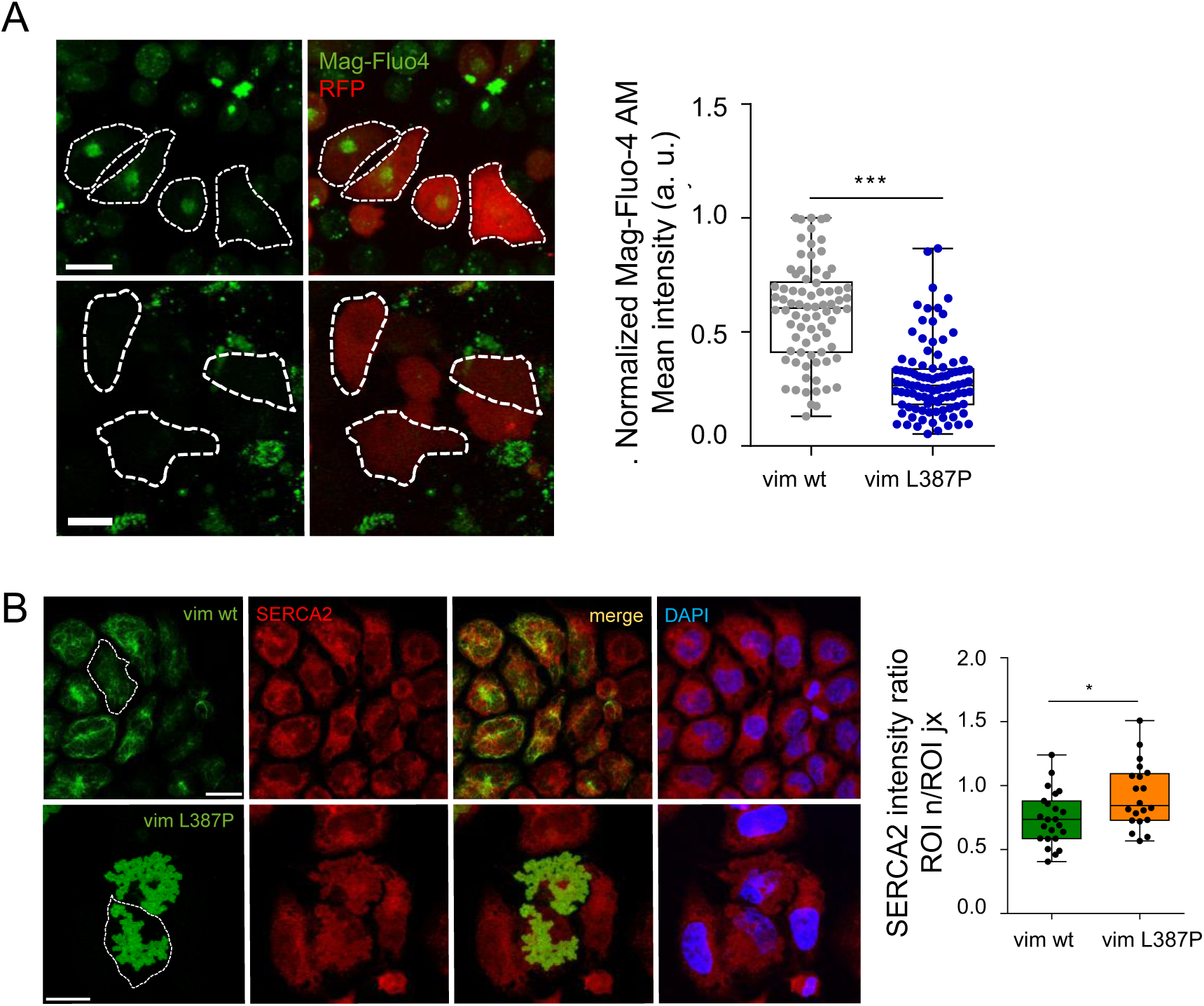
Vimentin L387P alters calcium levels and drives SERCA2 sequestration. (A) SW13/cl.2 cells transfected with RFP//vimentin wt or L387P were cultured in calcium and magnesium free HBSS medium containing 1 μM Mag-Fluo-4 AM for 10 minutes and then subjected to laser excitation at 488 nm to detect Mag-Fluo-4 AM fluorescence. Cell contours are indicated by dashed lines. Images represent single optical sections. Vim represents vimentin. Quantification of green signal intensity in RFP-positive cells (that is, cells expressing vimentin) is represented in the graph. (B) Distribution of SERCA2 in SW13/cl.2 cells transfected with pCMV6–vimentin wt or L387P and stained with DAPI. Confocal microscopy images represent total projections. SERCA2 redistribution, quantified as the ratio between the average fluorescence intensity of ROIs defined in the area corresponding to the nucleus (n) and at a juxtanuclear position (jx), is shown in the graph. (A) and (B) For all graphs, data from at least three independent experiments are presented as box plots, where boxes represent the interquartile range, the line indicates the median, and whiskers denote the full data range Statistical significance was assessed using the Student’s t-test. *p<0.05, ***p<0.001. Images shown are representative. Scale bars: 20 µm.

## Discussion

Vimentin plays a crucial role in preserving intracellular architecture. Beyond providing mechanical resilience, the vimentin filament network acts as a scaffold that positions and stabilizes multiple organelles, including the nuclei, mitochondria, lysosomes, the Golgi apparatus and the ER [5,8,10–12,59–62]. In the context of ER organization, vimentin interacts with the E3 ligase RNF26, a central ER quality-control factor, thereby promoting the perinuclear accumulation of ER membranes and supporting the structural integrity of ERAD hubs [12]. Additionally, lack of vimentin results in ER dispersion rather than perinuclear location [12,60]. Yet how disruption of vimentin architecture, rather than depletion, influences ER function remains poorly defined.

Missense mutations in the *VIM* gene provide a unique experimental lens to interrogate the functional relevance of vimentin filament integrity. The L387 residue within vimentin is highly conserved across the intermediate filament family [42,43], and mutations in this and the homologue residue in other IFs have been associated with severe pathologies [42,44–46]. With respect to vimentin, the L387P *de novo* substitution has been identified in a patient with a multisystem progeroid disorder [42]. This mutant failed to assemble into filaments in vimentin-deficient MEFs or MCF-7 cells, accumulating instead in prominent cytoplasmic aggregates. Co-expression with wt vimentin, however, largely restored filament network formation in most cells. Here, we report that the vimentin L387P mutant causes an aberrant vimentin perinuclear aggregate composed of interconnected tubular/cisternal-like structures in SW13cl.2 cells, in contrast to the extended and dynamic filament network formed by the wt protein. These aggregates frequently occupy the DAPI-defined nuclear area, and cells bearing the aberrant assembly exhibit amorphous nuclear shapes similar to those described in Hutchinson–Gilford progeria syndrome. [48,49].

It is worth noting that other pathological conditions also feature disrupted vimentin organization. For instance, several pathogenic missense mutations, such as E151K and Q208R, associated with congenital cataract, lead to cytoplasmic vimentin aggregates accompanied by impaired proteasome function [41,63]. In addition, posttranslational modifications induced by oxidative stress, a common hallmark of many diseases, can drive the reorganization of vimentin into abnormal structures and/or promote filament disassembly [6,13]. However, given the large size and characteristic morphology of the vimentin L387P aggregates, their pronounced perinuclear localization, and the fact that the ER is mainly distributed in this region, we focused our study on the vimentin L387P variant and its impact on ER organization. In SW13/cl.2 cells expressing vimentin wt, classical ER markers such as GRP78, GRP94, calreticulin, and CLIMP63 display the expected reticular and perinuclear pattern. In contrast, the vimentin L387P mutant induces a pronounced segregation of ER components: GRP78 and GRP94 accumulate within the vimentin L387P aggregate, whereas calreticulin and CLIMP63 are excluded and remain at the periphery of this abnormal structure. Because three of these four markers participate directly in ER quality control (ERQC), we next examined whether ERAD and ERAD-associated factors were similarly affected. We found that SEL1L, HRD1 and the 20S proteasome concentrate at the vimentin cisternal-like aggregate, whereas VCP/p97 and RNF26 are excluded. The sequestration of GRP78 and GRP94, two chaperones that assist nascent polypeptides and recognize exposed hydrophobic regions in unfolded proteins [36,52], together with their co-immunoprecipitation with vimentin, is consistent with the possibility that they engage misfolded L387P vimentin in an attempt to promote its correct folding.

A similar rationale may explain the accumulation of the 20S proteasome at the vimentin tubules. Aggresomes are known to recruit proteasomes, autophagosomes and lysosomes to promote aggregate clearance [64], and the 20S proteasomal subunit was found in ER-enriched membrane fractions after ER-stress [65]. In our experimental model, however, proteasome enrichment at the vimentin L387P aggregate does not result in effective degradation, as the mutant protein remains stable during a CHX chase, while vimentin wt levels clearly decline. Likewise, in SW13/cl.2 cells expressing vimentin L387P, ERAD substrates such as NHK and GFP-GluR1 accumulate instead of being cleared. These observations indicate that proteasomes positioned at the aggregate may be functionally impaired or rendered inaccessible to carry out their function, or, alternatively, that their presence reflects passive sequestration rather than the assembly of a productive degradative compartment. The mislocalization of core ERAD components further supports this interpretation. The SEL1L–HRD1 complex forms the central retrotranslocation module of mammalian ERAD, in which SEL1L stabilizes HRD1 and facilitates substrate engagement and ubiquitination [17,55,66]. In turn, VCP/p97 is required to extract ubiquitinated substrates into the cytoplasm for processing by the 26S proteasome, and can interact with HRD1 directly or indirectly through Derlin-1 and Derlin-2 [67,68]. The physical separation observed in this work of HRD1–SEL1L, which accumulate at the L387P mutant structure, from VCP/p97, which becomes predominantly nuclear, likely prevents productive retrotranslocation and explains the defective clearance of ERAD substrates. VCP/p97 is typically recruited to aggresomes [69], yet we observe the opposite localization pattern. This mismatch suggests that the L387P aggregate is structurally distinct from classical aggresomes and lacks the features necessary to support coordinated proteasomal degradation.

The exclusion of RNF26 from vimentin L387P aggregate further strengthens the view that ERQC architecture is disrupted. RNF26, through its C-terminal domain, binds vimentin and organizes a perinuclear ERQC compartment enriched in ERAD components, as shown by Cremer et al [12]. Through this interaction, vimentin functions as a scaffold that spatially integrates folding, quality control and the degradation machinery, in particular lysosomes. It could be speculated that the region of vimentin required for RNF26 binding, which remains uncharacterized, is masked or inaccessible within the vimentin L387P aggregate. This potential loss of RNF26–vimentin connectivity likely contributes to the failure to integrate ERAD modules with proteolytic compartments.

In this study, we evaluated key components of the ERQC/ERAD pathway across its major functional hubs [70]: chaperones such as GRP78 and calreticulin, responsible for substrate recognition; factors including SEL1L, HRD1 and VCP/p97, mediating substrate dislocation and retrotranslocation; E3 ligases such as RNF26 promoting ubiquitination; and proteasomal degradation through the catalytic proteasomal subunit 20S. At each of these levels, the examined proteins displayed profoundly altered localization in vimentin L387P expressing cells. Combined with the defective clearance of ERAD substrates, these findings indicate that the vimentin L387P mutant disrupts ER proteostasis.

Beyond proteostasis defects, our results point out that vimentin L387P impacts ER calcium handling. The ER is the primary intracellular calcium reservoir, and luminal Ca^2+^ is essential for the function of chaperones such as GRP78, GRP94, calreticulin and calnexin [71–73]. Therefore, perturbations in luminal Ca^2+^ compromise folding efficiency and can initiate ER stress [74]. Here, it was observed that in cells expressing vimentin L387P, there is a pronounced reduction in free Ca^2+^, while SERCA2, the ATP-dependent pump that restores the ER Ca^2+^ reservoir [31], loses its perinuclear enrichment and instead accumulates at the mutant vimentin aggregates. In MCF7 cells, inhibition of SERCA has been shown to disrupt ER Ca^2+^ homeostasis and severely compromise the folding environment, ultimately triggering apoptosis when sustained over time [75]. Given the increased cell death observed in vimentin L387P–expressing populations, it is plausible that the redistribution of SERCA2, together with the reduced ER Ca^2+^ levels reported here, contributes to the cytotoxicity associated with the L387P mutant.

The vimentin network plays a structural and functional role in lipid storage by forming a peridroplet cage that anchors lipid droplets through direct binding to perilipin. Loss of vimentin profoundly alters lipid droplet biogenesis and stability, leading to smaller adipocytes, reduced lipid stores, and impaired mobilization of neutral lipids [76]. Consistent with this, Cogné et al. reported that a patient affected by the progeroid syndrome caused by the vimentin L387P mutation showed partial lipodystrophy, and adipocytes expressing vimentin L387P presented lower perilipin expression with decreased number of lipid droplets and reduced amount of lipids within them [42].

Importantly, lipid droplets originate from and remain in continuity with the ER membrane, and alterations in ER membrane lipid content and organization provoke changes in lipid droplet composition [77]. The ER is the primary site for the synthesis of nearly all cellular membrane lipids, including phospholipids and cholesterol [78]. Thus, the lipid abnormalities reported in the patient carrying the vimentin L387P mutation are consistent with compromised ER function.

Perturbations in the ER phospholipidome can trigger chronic ER stress, alter the abundance of ER-resident proteins, and promote their premature degradation [79]. During membrane protein biogenesis, nascent proteins can transiently explore multiple topologies, and the surrounding lipid environment plays a decisive role in stabilizing their final orientation within the bilayer [80]. Consequently, changes in the ER lipid bilayer composition due to lipid bilayer stress can hamper the proper integration of transmembrane proteins and predispose them to early degradation [79]. It is therefore plausible that lipid imbalances arising from defective lipid synthesis or impaired lipid droplets biogenesis in vimentin L387P expressing cells contribute to the mislocalization or reduced stability of ERAD-associated transmembrane proteins such as HRD1, SEL1L, or SERCA2.

In conclusion, our findings show that the vimentin L387P mutation remodels the intermediate filament network into a perinuclear aggregate structure that traps a subset of ERQC components (i.e. GRP78, SEL1L, HRD1, and the 20S proteasomal core) while excluding others, such as calreticulin, VCP/p97, and RNF26. By spatially segregating ERAD components that must operate in close proximity to support substrate recognition, retrotranslocation and proteasomal delivery, the vimentin L387P aggregate disrupts the architectural continuity required for efficient clearance of misfolded proteins. These defects collectively compromise the ability of the ER to maintain protein homeostasis, and reduce intra-organelle Ca^2+^ reservoir capacity. Overall, our findings demonstrate that the integrity of the vimentin filament network is critical for preserving ER resident proteins distribution, ERAD function and calcium handling, and highlight pathogenic vimentin mutations as valuable experimental model for elucidating the cytoskeletal control of ER physiology.

## Abbreviations

CHX: cycloheximide
DAPI: 4′,6-diamidino-2-phenylindole
ER: endoplasmic reticulum
ERAD: endoplasmic reticulum associated degradation
ERQC: endoplasmic reticulum quality control
GFAP: glial fibrillary acidic protein
GluR1: glutamate receptor subunit 1
Mag-Fluo-4 AM: magnesiumsensitive fluorescent indicator acetoxymethyl ester
NHK: Null Hong Kong (misfolded α1-antitrypsin variant)
PLA: proximity ligation assay
SERCA2: sarco/endoplasmic reticulum Ca^2+^-ATPase 2.

## Author contributions

A.E.M. and E.S.L. performed the experiments and analyzed the data. A.E.M. and D.P.S. designed and supervised the study, and wrote the manuscript. D.P.S. secured and managed funding. All authors reviewed and approved the final version of the manuscript.

## Acknowledgements

We thank Dr. Kvido Strisovsky for providing plasmids, and the staff of the Flow Cytometry and Confocal Microscopy facilities at Centro de Investigaciones Biológicas Margarita Salas for their technical assistance and advice.

