## Supplementary material for "The pathogenic mutant vimentin L387P disrupts endoplasmic reticulum organization and proteostasis"

Running title: Vimentin L387P drives ER disorganization and dysfunction

This file includes:

Figs. S1 to S2

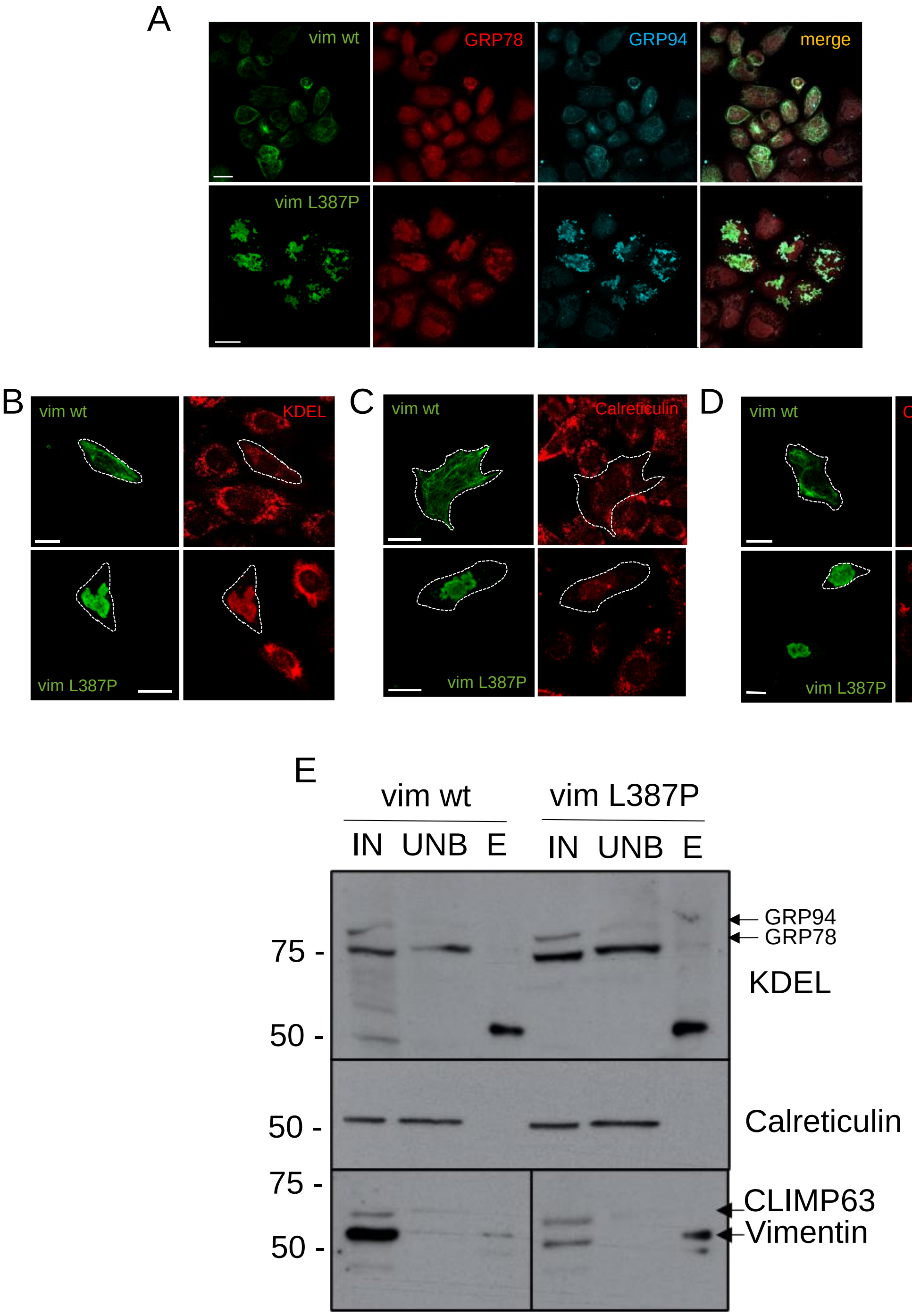


**Supplementary Figure S1. ER marker distribution and vimentin-associated chaperone pull-down in vimentin wt and L387P-expressing cells.** (A) Immunofluorescence analysis of vimentin (vim), GRP78, and GRP94 in SW13/cl.2 cells transiently transfected with pCMV6–vimentin wt or L387P. (B–D) MEF *Vim(–/–)* cells transfected with RFP//vimentin wt or L387P were stained for (B) GRP78/GRP94 (anti-KDEL antibody), (C) calreticulin, or (D) CLIMP63, together with vimentin. Cell contours are indicated by dashed lines. Confocal microscopy images represent total projections. Representative images from three independent experiments are shown. Scale bars: 20 µm. (E) Vimentin immunoprecipitation from total lysates of SW13/cl.2 cells expressing vimentin wt or L387P was performed using the anti-vimentin V9-AC antibody. Immunoprecipitates were analyzed by western blot and probed for GRP78/GRP94, calreticulin, CLIMP63, and vimentin. The position of the bands of interest is indicated by arrows.

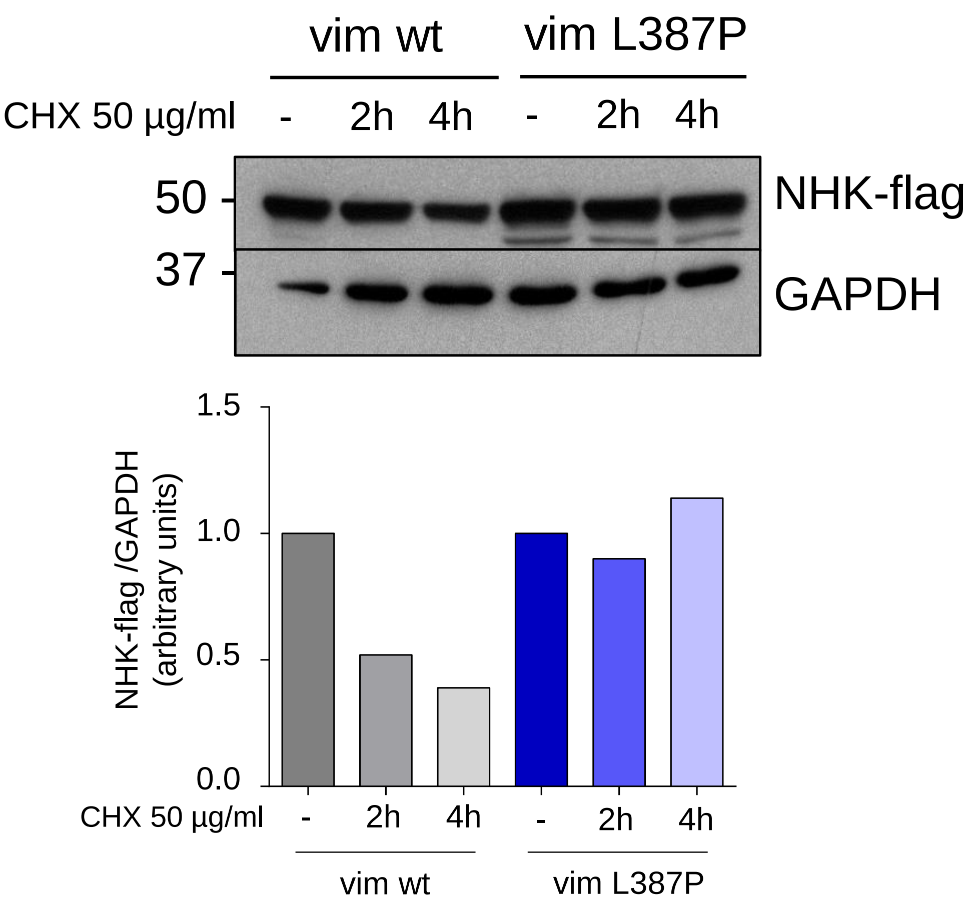


**Supplementary Figure S2. ERAD substrate turnover in MEF *Vim(-/-)* expressing vimentin wt or L387P.** Cycloheximide (CHX) chase assay of NHK-flag degradation in MEF *Vim(-/-)* transiently co-transfected with RFP//vimentin wt or L387P and NHK-flag. Vim refers to vimentin. Cells were treated with 50 µg/mL CHX for 0, 2, or 4 h. Lysates were analyzed by Western blot using anti-flag to detect NHK-flag. GAPDH was used as a loading control. The graph depicts quantification of NHK-flag levels normalized to GAPDH and expressed relative to the control condition from an assay selected as representative. Images are representative from two independent experiments.
